# Location analysis of presynaptically active and silent synapses in single-cultured hippocampal neurons

**DOI:** 10.1101/2023.12.12.571354

**Authors:** Otoya Kitaoka, Kohei Oyabu, Kaori Kubota, Takuya Watanabe, Satoru Kondo, Teppei Matsui, Shutaro Katsurabayashi, Katsunori Iwasaki

**Author notes:** Correspondence should be addressed to: Prof. Shutaro Katsurabayashi, Ph.D., Department of Neuropharmacology, Faculty of Pharmaceutical Sciences, Fukuoka University, 8-19-1 Nanakuma, Jonan-ku, Fukuoka 814-0180, Japan., (ext. 6634).

## Abstract

A morphologically present but non-functioning synapse is termed a silent synapse. Silent synapses are categorized into “postsynaptically silent synapses,” where AMPA receptors are either absent or non-functional, and “presynaptically silent synapses,” where neurotransmitters cannot be released from nerve terminals. The presence of presynaptically silent synapses remains enigmatic, and their physiological significance is highly intriguing. In this study, we examined the distribution and developmental changes of presynaptically active and silent synapses in individual neurons. Our findings show a gradual increase in the number of excitatory synapses, along with a corresponding decrease in the percentage of presynaptically silent synapses during neuronal development. To pinpoint the distribution of presynaptically active and silent synapses, i.e., their positional information, we enhanced the traditional Sholl analysis and introduced a novel method termed “donut analysis.” Our results indicate that the distribution of presynaptically silent synapses within a single neuron does not exhibit a distinct pattern during synapse development in different areas. However, irrespective of neuronal development, the proportion of presynaptically silent synapses tends to rise as the projection site moves farther from the cell body, suggesting that synapses near the cell body may exhibit higher synaptic transmission efficiency. This study represents the first observation of changes in the distribution of presynaptically active and silent synapses within a single neuron. Additionally, we propose that donut analysis can serve as a valuable analytical tool for evaluating synaptic positional information.

**Scope statement:** A morphologically present but non-functioning synapse is termed a silent synapse. The presence of presynaptically silent synapses remains enigmatic, and their physiological significance is highly intriguing. This study focused on the distribution and developmental changes of presynaptically active and silent synapses in individual neurons. To pinpoint the distribution of presynaptically active and silent synapses, we enhanced the traditional Sholl analysis and introduced a novel method termed “donut analysis.” We found that the distribution of presynaptically silent synapses within a single neuron does not exhibit a distinct pattern during synapse development in different areas. However, irrespective of neuronal development, the proportion of presynaptically silent synapses tends to rise as the projection site moves farther from the cell body.

This is a new paper that applies “Sholl analysis,” a method invented 70 years ago that is now the gold standard of morphological analysis of the single neuron.

This study represents the first observation of changes in the distribution of presynaptically active and silent synapses within a single neuron. Additionally, we propose that donut analysis can serve as a valuable analytical tool for evaluating synaptic positional information for the design of “synaptic maps” in neural circuits.

## 1. Introduction

An excitatory synapse releases glutamate as a neurotransmitter, and it is referred to as an active synapse. Conversely, a synapse that maintains its synaptic structure but fails to transmit neuronal information is known as a silent synapse (1). Silent synapses can become inactive for one of two reasons: (I) the absence or impairment of receptor function in the post-synaptic membrane or (II) the loss of synaptic exocytotic function in the nerve terminal.

Regarding reason(I), the relationship between two types of glutamate receptors, namely the N-methyl-D-aspartate (NMDA) receptor and the α-amino-3-hydroxy-5-methyl-4-isoxazolepropionic acid (AMPA) receptor, is speculated as follows: In the immature brain shortly after birth, NMDA receptors are expressed but blocked by Mg^2+^. Even when neurotransmitters are released and glutamate is received, NMDA receptors remain inactive, and neuronal information is not transmitted to subsequent neurons (2,3). However, as the brain matures, AMPA receptors appear near NMDA receptors. When released glutamate binds to AMPA receptors, depolarization occurs, removing the magnesium block at the NMDA receptor, leading to the activation of the synapse (4). Consequently, the nerve cell becomes more excited and transmits information to the next cell. In contrast to the mechanism described above, the physiological significance of reason (II) has not been fully elucidated (5).

*In vitro* conditions allowed for electrophysiological and morphological analyses, revealing notable distinctions in neuronal synaptogenesis (6,7,8,9). Building upon widely accepted concepts concerning presynaptic synaptogenesis, this study delves into presynaptic synaptogenesis within cultures spanning 1 week, 2 weeks, and 3 weeks. Our investigation also places particular emphasis on the distance from the cell body in relation to the proportion of presynaptically active and silent synapses projecting to dendrites.

## 2. Materials and methods

### 2.1 Animal ethics

All animal care procedures followed the rules of the Fukuoka University Experimental Animal Welfare Committee (equivalent to NIH guidelines). The experiment was strictly conducted after the Committee’s approval of the experimental plan. Cultured cells were obtained by decapitating newborn mice, and efforts were made to minimize distress.

All experiments were performed in compliance with the ARRIVE guidelines. Experiments were performed blind.

### 2.2 Experimental animals

Timed-pregnant Jcl:ICR mice (Catalog ID: Jcl:ICR, CLEA Japan, Inc., Tokyo, Japan) were purchased at gestational day 15 from the Kyudo Company (Tosu, Japan). Fifteen to seventeen-week-old pregnant Jcl:ICR mice were used. The pregnant mice were housed in plastic cages in an environment with a room temperature of 23±2°C, a humidity of 60±2%, and a 12-h light–dark cycle (lights on at 7:00 AM, lights off at 7:00 PM). Food (CLEA Rodent Diet, CE-2, CLEA Japan, Inc., Tokyo, Japan) and water were provided *ad libitum*. The body weights of pregnant mice were not recorded.

Experimental animals were handled in accordance with the animal ethics regulations of the Fukuoka University Animal Care and Use Committee (Approval No. 2112094 and 2311081).

### 2.3 Autaptic culture preparation

A sample in which a single neuron is cultured on a dot-like layer of astrocytes is referred to as an autaptic culture (11). The autaptic culture preparations were conducted in accordance with previous reports (11,12). To provide a brief overview, postnatal day 0–1 neonatal mice were used, and their brains were extracted and immersed in Hank’s Balanced Saline Solution (Invitrogen, Cat. # 084-08345) cooled to 4°C. In this state, the cerebral cortices on both sides were excised under a microscope, and cerebral cortical cells were isolated via trypsinization. The isolated cells were then cultured in 75-cm^2^ culture flasks (Corning Inc., NY, USA). After 2 weeks, the culture flask was gently tapped multiple times to remove non-astrocytic cells. Subsequently, the astrocytes that remained in close contact with the bottom of the culture flask were detached using trypsinization. These cells were replated at a density of 6,000 cells/cm^2^ per well onto 22-mm round coverslips (thickness No. 1; Matsunami, Osaka, Japan) within 6-well plates (TPP, Switzerland).

To cultivate the seeded astrocytes in dot shapes, a mixture of collagen (final concentration 1.0 mg/mL; BD Biosciences, San Jose, CA, USA) and poly-D-lysine (final concentration 0.25 mg/mL; Sigma-Aldrich, St Louis, MO, USA) was prepared. Subsequently, 300-μm square dots were stamped onto a round cover glass pre-coated with 0.5% agarose. This stamp design was an original development. One week after seeding the astrocytes, it was confirmed that the astrocytes had successfully formed dot-shaped cultures.

Next, brains were excised from neonatal ICR mice on days 0–1 after birth and immersed in Hank’s Balanced Saline Solution cooled to 4°C. In this state, the hippocampal CA3–CA1 region was dissected under a microscope. Finally, hippocampal neurons were isolated through treatment in Dulbecco’s modified Eagle’s medium (Invitrogen) containing 2 U/ml of papain (Worthington, Cat. # PAP) at 37°C for 1 hour. The isolated hippocampal neurons were then seeded at a density of 1,500 cells/cm^2^ per well and cultured in a 37°C, 5% CO_2_ incubator. Data from three groups (1 week, 2 weeks, and 3 weeks *in vitro*, respectively) were obtained from the same sister cultures (15 cultures in total).

### 2.4 FM1-43FX dye staining

Presynaptic terminals that actively release neurotransmitters, referred to as active synapses, were visualized using N-(3-triethylammoniumpropyl)-4-(4-(dibutyl amino) styryl) pyridinium dibromide (FM1-43FX, a fixable analog of FM1-43 membrane stain, Thermo Fisher Scientific, Waltham, MA, USA). To stain the presynaptically active synapses of autaptic cultured neurons, we followed the method of Moulder et al. (5,13). In brief, we dissolved 10 μM FM1-43FX in a high potassium (45 mM) extracellular solution containing the NMDA receptor inhibitor (2R)-amino-5-phosphonovaleric acid (APV, 25 µM, Sigma-Aldrich, St Louis, MO, USA) and the AMPA receptor inhibitor 6-cyano-7-nitroquinoxaline-2,3-dione (CNQX, 10 µM, Sigma Aldrich, St Louis, MO, USA). This solution was applied to the autaptic culture neurons for 2 min. Subsequently, the cells were washed three times for 2 min each with a standard extracellular solution containing 1 μM tetrodotoxin (TTX), a sodium channel blocker.

Following the staining procedure, autaptic culture neurons were fixed using a 4% paraformaldehyde solution in phosphate-buffered saline (PBS) for 10 minutes. To minimize the loss of FM1-43FX signals, such as photobleaching due to ambient light exposure, the images were captured promptly after fixing the neurons. We acquired sixteen-bit images using an all-in-one fluorescence microscope (BZ-X810, KEYENCE, Osaka, Japan) with a 20× objective lens (Plan Apochromat, numerical aperture 0.75), or an sCMOS camera (pco.edge 4.2, pco, Kelheim, Germany) mounted on an inverted microscope (Eclipse-TiE, Nikon, Tokyo, Japan) equipped with a 40× objective lens (Plan Apoλ, numerical aperture 0.95). In the case of using the inverted microscope, FM1-43FX was excited using a white LED (Lambda HPX, Sutter Instruments, Novato, CA, USA) at 100% maximum intensity and imaged using a filter cube (470/40-nm excitation, 500-nm dichroic long-pass, 535/50-nm emission). In each sample, ten images were captured with an exposure time of 300 ms per image, averaged, and utilized for analysis based on the average pixel intensity.

### 2.5 Immunostaining

Autaptic culture preparations underwent immunostaining based on the method established by Moulder et al. (5,13). After capturing FM1-43FX images, autaptic culture neurons were incubated in a microscope chamber with PBS containing 5% normal goat serum and 0.1% Triton X-100 (Sigma Aldrich, St Louis, MO, USA) for 30 minutes. Following the Triton X-100 blocking step, the decolorization of FM1-43FX was visually confirmed (data not shown). Primary antibodies were subsequently applied for 3 h at the following dilutions: anti-microtubule-associated protein 2 (MAP 2) at 1:1,000 (guinea pig polyclonal, antiserum, Synaptic Systems, Göttingen, Germany) and anti-vesicular glutamate transporter 1 (anti-vGLUT1) at 1:2,000 (rabbit polyclonal, affinity-purified, Synaptic Systems, Göttingen, Germany). Secondary antibodies were applied using Alexa Fluor 488 or 594 (Thermo Fisher Scientific, Waltham, Mass., USA) at a dilution of 1:400 for 30 min. Since FM1-43FX can be completely removed by Triton X-100 blocking (5,12), the excitation light (480 nm) used for fluorescence observation of FM1-43FX was also employed for fluorescence excitation of Alexa Fluor 488.

Imaging of autaptic culture preparations was performed using an all-in-one fluorescence microscope (BZ-X810, KEYENCE, Osaka, Japan) with a 20× objective lens (Plan Apochromat, numerical aperture 0.75), or an sCMOS camera (pco.edge 4.2, pco, Kelheim, Germany) mounted on an inverted microscope (Eclipse-TiE, Nikon, Tokyo, Japan) equipped with a 40× objective lens (Plan Apoλ, numerical aperture 0.95). Similar to FM1-43FX imaging, ten images were captured per sample, and these images were subsequently normalized to obtain the average intensity for analysis.

### 2.6 Qualification of synaptic puncta

To identify the vGLUT1 puncta, we employed ImageJ software (version 1.46j; Rasband, W.S., ImageJ, U. S. National Institutes of Health, Bethesda, Maryland, USA, https://imagej.nih.gov/ij/, 1997-2016). We subtracted the original images from an image filtered with a Gaussian blur of the duplicated original image. For detailed procedures, please refer to Iwabuchi et al. (14). The subtracted images were then subjected to binarization using a threshold set at the top of 0.01% of the cumulative intensity of the background area in the vGLUT1 image. Subsequently, we detected the number of puncta overlaid with MAP2 images, applying a size threshold of ≥ 5 pixels.

### 2.7 Qualification of presynaptically silent synapses

The FM1-43FX puncta were identified in a manner similar to that employed for vGLUT1 puncta. We overlaid images of FM1-43FX with images of vGLUT1 and MAP2 to identify presynaptically silent synapses. Utilizing ImageJ, we defined the region of interest (ROI) of vGLUT1 that was not stained with FM1-43FX as a presynaptically silent synapse. For a more comprehensive understanding of the analysis of silent synapses, please refer to our previous study (12).

### 2.8 Donut analysis

The Sholl analysis plugin (15) within Image J was employed to examine the projection positions of presynaptically silent synapses. The methodology for this analysis is outlined as follows: Initially, a minimum circle with a diameter of 20 µm was delineated around the cell body. Subsequently, concentric circles were drawn to encompass the entire MAP2 image, including three additional concentric circles positioned between the maximum circle and the central minimum circle (Fig. 2A). The region inside the minimal circle was designated as area 1, the region between the outer edge of the minimum circle and the subsequent concentric circle was labeled as area 2, the area extending to the next concentric circle was designated as area 3, the region encompassing the following concentric circle was defined as area 4, and the outermost region was identified as area 5 (Fig. 2B). By categorizing the synapses present within each of these donut-shaped areas, it becomes possible to quantify the positional information of the synapses. We coined this analytical method “donut analysis” due to the donut-shaped nature of the areas under investigation.

### 2.9 Statistics

Data are expressed as mean ± SEM. Statistical tests were conducted using Matlab Statistics Toolbox (MathWorks, Natick, M.A.). Developmental changes of total number of synapses or total proportion of synapses were evaluated using Pearson correlation coefficients between the culture period and number of synapses or the culture period and proportion of synapses, respectively (Fig. 1B-E). Developmental changes of numbers and proportions of synapses across areas were evaluated using two-way analysis of variance followed by Tukey’s HSD test (Fig. 2C-E). Differences were considered significant when *p* < 0.05.

**Figure 1.**
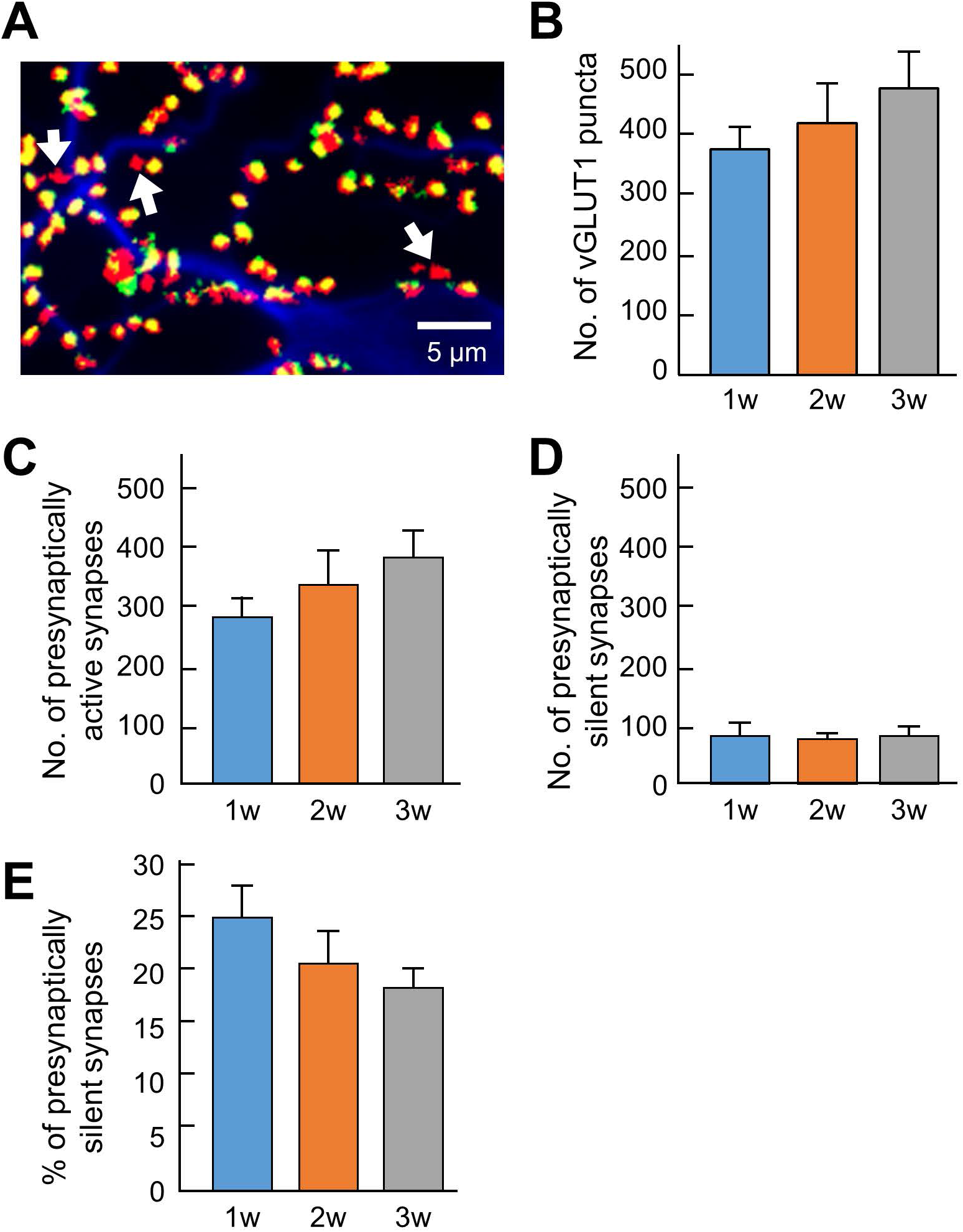
Changes in the number of synapses during development. (A) Typical fluorescence image. In terms of pseudocolors, vGLUT1 is labeled in red, FM1-43FX is in green, and MAP2 is in blue. Therefore, when the green staining of FM1-43FX and the red staining of vGLUT1 overlap, presynaptically active synapses appear yellow. Conversely, presynaptically silent synapses lack the green label and are denoted as red-only puncta by vGLUT1 (as indicated by the arrows in Fig. 1A). (B) Quantification of the number of vGLUT1-positive synapses (blue bar: 1 w: *n* = 33, orange bar: 2 w: *n* = 26, gray bar: 3 w: *n* = 32). (C) Quantification of the number of presynaptically active synapses (blue bar: 1 w: *n* = 33, orange bar: 2 w: *n* = 26, gray bar: 3 w: *n* = 32). Data were obtained from the same neuron as in Figure 1B. (D) Quantitation of the number of presynaptically silent synapses (blue bar: 1 w: *n* = 33, orange bar: 2 w: *n* = 26, gray bar: 3 w: *n* = 32). Data were obtained from the same neuron as in Figure 1B. (E) Percentage of presynaptically silent synapse numbers (blue bar: 1 w: *n* = 33, orange bar: 2 w: *n* = 26, gray bar: 3 w: *n* = 32). Data were obtained from the same neuron as in Figure 1B.

**Figure 2.**
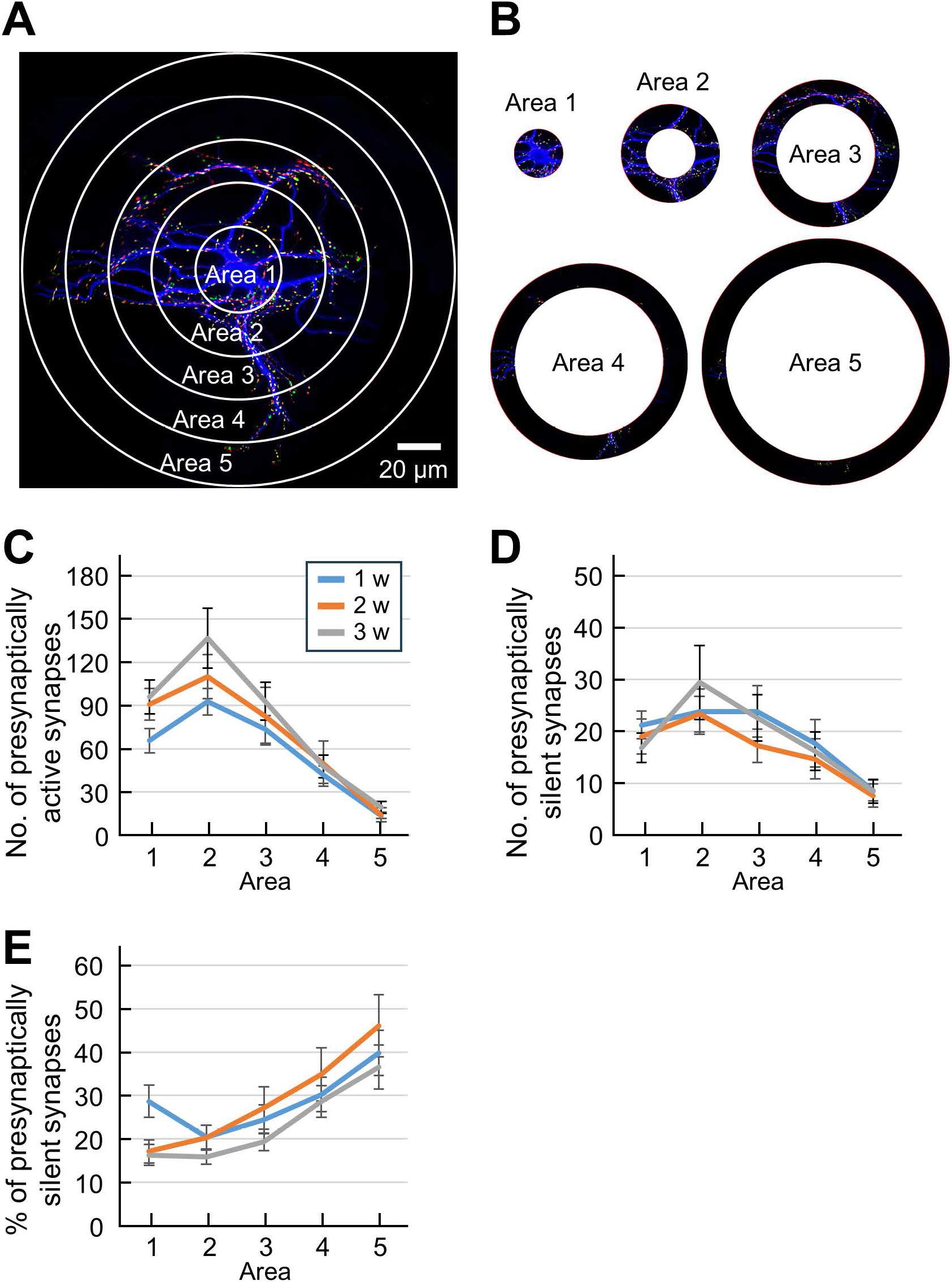
Quantification of synaptic location information by donut analysis. (A) Fluorescence image of an autaptic culture preparation before sectioning into donuts. The inside of the minimal circle is area 1, the area from the outside of the minimum circle to the next concentric circle is area 2, the area to the next concentric circle is area3, the area to the next concentric circle is area 4, and the outermost area is area 5. (B) Sectioned donut area cut away. (C) Quantification of the number of presynaptically active synapses in each area (blue line: 1 w: *n* = 33, orange line: 2 w: *n* = 26, gray line: 3 w: *n* = 32). The horizontal axis indicates the area number. Data were obtained from the same neuron as in Figure 1B. (D) Quantification of the number of presynaptically silent synapses in each area (blue line: 1 w: *n* = 33, orange line: 2 w: *n* = 26, gray line: 3 w: *n* = 32). The horizontal axis indicates the area number. Data were obtained from the same neuron as in Figure 1B. (E) Percentage of presynaptically silent synapse numbers per area (blue line: 1 w: *n* = 33, orange line: 2 w: *n* = 26, gray line: 3 w: *n* = 32). The horizontal axis indicates the area number. Data were obtained from the same neuron as in Figure 1B.

## 3. Results

### 3.1 Quantification of presynaptically active and presynaptically silent synapses

FM1-43FX is internalized into the presynaptic terminals through the process of synaptic vesicle endocytosis. Consequently, we used fluorescent puncta labeling to identify presynaptically active synapses capable of neurotransmitter exocytosis. Figure 1A illustrates a representative fluorescence image in which vGLUT1 is pseudocolored in red, FM1-43FX in green, and MAP2 in blue. In the case of presynaptically active synapses, the overlap of the green FM1-43FX and red vGLUT1 stains results in a yellow appearance, indicating the presence of a presynaptically active synapse. Conversely, presynaptically silent synapses, which fail to uptake FM1-43FX, are not marked in green but appear solely as red puncta by vGLUT1 (indicated by arrows in Fig. 1A). Essentially, the red fluorescent puncta denote presynaptically silent synapses, which are excitatory synapses that do not release glutamate through exocytosis.

First, we quantified the number of synapses positive for the vGLUT1 antibody. Although not statistically significant, the results showed a gradual increase in the number of excitatory synapses as neurons developed (Fig. 1B, 1 w: 381.58±36.63, 2 w: 426.58±62.55, 3 w: 486.56±57.11; R = 0.16, *p* = 0.14). It is important to note that the vGLUT1 puncta in this result represents both presynaptically active and presynaptically silent synapses.

To specifically quantify presynaptically active synapses, we counted the number of synapses labeled with FM1-43FX among vGLUT1-positive synapses. The results indicated a gradual increase in the number of presynaptically active synapses as neurons developed (Fig. 1C, 1 w: 287.09±29.42, 2 w:347.65±54.92, 3 w: 393.13±46.25), though the correlation was not significant (R = 0.19, *p* = 0.075).

Next, we quantified presynaptically silent synapses among vGLUT1-positive synapses, identified as those where FM1-43FX was not labeled (Fig. 1D). The results revealed no change in the number of presynaptically silent synapses with neuronal development (Fig. 1D, 1 w: 94.48±16.76, 2 w: 81.88±12.87, 3 w: 93.44±14.93; R = - 0.0059, *p* = 0.96). Based on these findings, we calculated the ratio of presynaptically silent synapses (Fig. 1E) and observed that the proportion significantly decreased with neuronal development (Fig. 1E, 1 w: 25.04±2.75%, 2 w: 20.91±2.73%, 3 w: 18.45±1.61%; R = -0.2097, *p* < 0.05).

### 3.2 Location analysis of synapses using donut analysis

Upon observing the image in Fig. 1A, we noted that the projection positions of presynaptically active and silent synapses onto the dendrites exhibited uneven distribution. Consequently, we endeavored to quantify the positional information of these synapses within a single neuron. To conduct this positional analysis of synaptic puncta, we employed the conventionally known Sholl analysis (15). Sholl analysis is a widely used and straightforward method for quantifying the branching patterns of dendrites and axons. In this study, concentric circles were drawn around the neuron’s cell body (Fig. 2A), and these concentric circles were then divided into donut-shaped areas (Fig. 2B). Synapses were tallied within each of these donut-shaped regions, enabling precise quantification of the synapse distribution.

The number of presynaptically active synapses in each area exhibited a peak in area 2 for all three groups (Fig. 2C). Significant differences were found between area 2 and area 5 for all three groups and between area 2 and area 4 for 2 and 3 weeks (*p* < 0.05). In general, the number of presynaptically active synapses decreased as the distance from the cell body increased (Fig. 2C) (1 w, R = -0.42, *p* < 0.05; 2 w, R = - 0.38, *p* < 0.05; 3 w, R = -0.42, *p* < 0.05). Similarly, the number of presynaptically silent synapses in each area exhibited a relative decrease as the distance from the cell body increased (Fig. 2D) (1 w, R = -0.19, *p* < 0.05; 2 w, R = -0.26, *p* < 0.05; 3 w, R = -0.17, *p* < 0.05). Based on these findings, we calculated the percentage of presynaptically silent synapses for each area (Fig. 2E). Intriguingly, the proportions of presynaptically silent synapses near the cell body (area 1 and area 2) and that of the most distal part of the cell body (area 5) were significantly different for 2 and 3 weeks (p < 0.05) but not for 1 week, indicating developmental change of the distribution of presynaptically silent synapses. Additionally, the location analysis revealed a statistically significant trend of an increasingly higher proportion of presynaptically silent synapses as the distance from the cell body increased (Fig. 2E) (1 w, R = 0.20, *p* < 0.05; 2 w, R = 0.37, *p* < 0.05; 3 w, R = 0.38, *p* < 0.05).

## 4. Discussion

Compared to a previous study (16), where the percentage of active synapses was approximately 66%, our experiments revealed percentages ranging from about 75-80% (Fig. 1E). Clearly, these disparities can be attributed to extrinsic factors such as the culture conditions of the neurons, though differences in experimental methods cannot be discounted. In the previous study (16), FM dye staining was conducted using action potential trains and treated with a high potassium solution for FM dye-destaining. In our present experiment, we employed robust stimuli, including a high-potassium solution. It is plausible that such a potent stimulus may have “awakened” dormant presynaptic synapses. Therefore, measuring functional active presynapses through electrical stimulation is an avenue for future investigation.

The increase in the number of excitatory synapses with neuronal development is a well-documented phenomenon (17,18), and the findings of our study align with this observation (Fig. 1B). Turning attention to the percentage of presynaptically silent synapses within each compartment, we observed that approximately 20% remained silent in areas 2–3, while approximately 40% were silent in area 5, indicating that silent synapses tend to form at a greater distance from the cell body. This suggests a trend toward an increased presence of excitatory active synapses proximal to the soma (Fig. 2C). Synapses in close proximity to the cell body were posited to be more active than those distal to the cell body during neuronal development, with two potential explanations. Firstly, as synapses develop, they undergo synaptic pruning, a process where axons reshape neuron dendrites and synapses, eliminating unnecessary synapses during brain development (19,20,21,22). Synapses were initially formed across the entire dendritic structure in immature neurons, with presynaptically silent synapses near the soma being pruned during development. Secondly, presynaptic synapses situated far from the soma may undergo slower development. Activation of presynaptically silent synapses in immature neurons is believed to be contingent on the PKA signaling pathway (23,24). Consequently, synaptic function may evolve in tandem with neuronal development, especially in the vicinity of the cell body, where second messengers exert influence and activate previously inactive synapses.

In this study, we did not observe significant changes in the number of presynaptically silent synapses during development (Fig. 1D). It is worth noting that changes in the number of postsynaptically silent synapses during development have been reported (25). For instance, in the neonatal rat visual cortex, many silent synapses exist in layer VI pyramidal neurons, and the number of active synapses increases with growth, similar to the hippocampus. On the other hand, layer II/III pyramidal cells have many active synapses at birth, and silent synapses increase with growth, followed by a return to active synapses (25). Thus, the patterns of developmental post-synaptic expression appear to vary by brain region. It remains unclear whether the results of this study are specific to glutamatergic neurons in the hippocampus or if similar patterns are observed in other brain regions.

While no distinct changes were observed in the ratio of silent synapses in each area during neuronal development, an interesting finding was that, regardless of neuronal maturation, the proportion of presynaptically silent synapses was lower in the proximal region compared to the distal region of the cell body. Although the physiological significance of changes in the rate of presynaptically silent synapse formation during neuronal development remains unknown, it may contribute to the establishment of functional neural circuits.

Presynaptically silent synapses, despite being structurally mature, are believed to lack neurotransmitter release due to the inability of synaptic vesicles to exocytose (24). Several proteins, such as Munc13, RIM, CAST, and bassoon, are involved in neurotransmitter exocytosis from nerve terminals (26,27). Among these, Rim1 and Munc13-1 have been reported to decrease in expression after the induction of silent synapses following depolarization induction in hippocampal neurons (28). It remains unclear whether such presynaptic proteins associated with exocytosis from nerve terminals are more highly expressed at synapses projecting closer to the cell body. Renger et al. (10) revealed that functional synaptic vesicle turnover follows the localization of synapsin I with a 1-2 day delay. However, the primary factor distinguishing early-stage synaptogenesis from silent synapses remains unknown.

The NMDA receptor NR2B subunit has been reported to be replaced by NR2A during neuronal maturation, resulting in decreased exocytosis (29). Based on these results, presynaptically silent synapses may possibly increase due to premature maturation of synapses closer to the distal cell body as neurons develop, along with an increase in the NR2A subunit. To verify this hypothesis, a qualitative determination of the expression position of the NR2A subunit is necessary. However, the regulation of expression and location of presynaptically silent synapses during development remains unclear, necessitating further research in the future.

It may be challenging to ensure that donut analysis accurately distinguishes distal from proximal synapses. For instance, axons and dendrites can freely extend within the astrocytic dot area, and the possibility that proximal concentric rings capture distal synapses cannot be dismissed. However, this concern applies similarly to conventional Sholl analysis. Nonetheless, we trust that the findings of this study will offer insights into unraveling the mechanism of synapse development, including its unknown potential, significance, and functions.

Above all, we aspire for this innovative technique, “donut analysis,” to become a valuable method for analyzing the positional information of synapses within a single neuron.

## 5. Data Availability

The data that support the findings of this study are available from the corresponding author upon reasonable request.

## 6. Author Contributions Statement

O.K., K.O., and S.K. (Shutaro Katsurabayashi) performed experiments and analyzed data; K.K., T.W., S.K. (Shutaro Katsurabayashi), and K.I. conceived the study; S.K. (Satoru Kondo), and T.M., K.I interpreted the data; K.O. and S.K. (Shutaro Katsurabayashi) wrote the manuscript with input from all authors. All authors reviewed the manuscript.

### 7.2 Ethical publication statement

We confirm that we have read the Journal’s position on issues involved in ethical publication and affirm that this report is consistent with those guidelines.

## 7.3 Acknowledgments

We thank the following members for supporting the preliminary experiments and analyzing the data: Miharu Koyama, Mika Suenaga, and Shuya Tsuneoka. This work was supported by a KAKENHI Grant-in-Aid for Transformative Research Areas (B) “Multicellular Neurobiocomputing” (21H05165), and a KAKENHI Grant-in-Aid for Scientific Research (B) (20H04506, 23H02805, and 23H02884, and 23H02572) from the Japan Society for the Promotion of Science.

